# Degradation of recalcitrant polyurethane and xenobiotic additives by a selected landfill microbial community and its biodegradative potential revealed by proximity ligation-based metagenomic analysis

**DOI:** 10.1101/760637

**Authors:** Itzel Gaytán, Ayixon Sánchez-Reyes, Manuel Burelo, Martín Vargas-Suárez, Ivan Liachko, Maximilian Press, Shawn Sullivan, M. Javier Cruz-Gómez, Herminia Loza-Tavera

**Affiliations:** Departamento de Bioquímica, Facultad de Química, Universidad Nacional Autónoma de México, Ave. Universidad 3000, Col. UNAM, Ciudad de México, 04510, MÉXICO.; Departamento de Química Analítica, Facultad de Química, Universidad Nacional Autónoma de México, Ave. Universidad 3000, Col. UNAM, Ciudad de México, 04510, MÉXICO.; Phase Genomics Inc, 1617 8th Ave N, Seattle, WA 98109, USA.; Departamento de Ingeniería Química, Facultad de Química, Universidad Nacional Autónoma de México, Ave. Universidad 3000, Col. UNAM, Ciudad de México, 04510, MÉXICO.

## Abstract

Polyurethanes (PU) are the sixth more produced plastics with around 19-million tons/year, but since they are not recyclable they are burned or landfilled, generating ecological damage. To elucidate the mechanisms that landfill microbial communities perform to attack recalcitrant PU plastic, we studied the BP8 community selected by its capability to grow in a water PU dispersion (WPUD) that contains a polyether-polyurethane-acrylate (PE-PU-A) copolymer and xenobiotic additives (N-methyl 2-pyrrolidone, isopropanol and glycol ethers), and performed a proximity ligation-based metagenomic analysis for revealing the community structure and potential biodegradative capacity. Additives were consumed early whereas the copolymer was cleaved throughout the 25-days incubation. BP8 metagenomic deconvolution reconstructed five genomes, three of them from novel species. Genes encoding enzymes for additives biodegradation were predicted. The chemical and physical analysis of the biodegradation process, and the identified biodegradation products show that BP8 cleaves esters, aromatic urethanes, C-C and ether groups by hydrolytic and oxidative mechanisms. The metagenomic analysis allowed to predicting comprehensive metabolic pathways and enzymes that explain the observed PU biodegradation. This is the first study revealing the metabolic potential of a landfill microbial community that thrives within a WPUD system and shows potential for bioremediation of polyurethane- and xenobiotic additives-contaminated sites.

## INTRODUCTION

Plastic pollution represents a pervasive anthropogenic threat for the survival of natural ecosystems. Worldwide, plastics have become so abundant that they have been proposed as geological markers for the Anthropocene era [1]. In 2017, 348 million tons of plastics were manufactured [2] and their production keeps increasing. Polyurethanes (PU) are versatile plastics produced as thermoplastics, thermosets, coatings, adhesives, sealants and elastomers that are incorporated into our daily life in building insulation, refrigerators and freezers, furniture and bedding, footwear, automotive, coatings, adhesives, and others. PU has been ranked as the sixth most used polymer worldwide with a production of 18 million tons in 2016 [3, 4]. The extensive utilization of PU generates wastes that are mainly disposed in municipal landfills where, because of its structural complexity will remain as polymeric structures for decades, or are burned generating toxic substances that negatively impact human health and ecosystems [3]. Furthermore, some PU such as polyether (PE)-PU are more recalcitrant than others, and additionally, some polyurethane-based liquid formulations contain additives that include secondary alcohols and glycol ethers that function as solvents or coalescing agents. Glycol ethers enter the environment in substantial quantities, are toxic for many microbial species [5–7] and represent a potential hazard for human health [8].

Over the last three decades, several research groups have isolated microorganisms with capacity to attack PU [9–15] and degrade xenobiotic additives [7, 9, 16, 17], and the abilities from several fungal and bacterial communities have been assessed in compost, soil, or liquid cultures [18–21] and in different activated sludges [22–25]. However, PU biodegradation is still a challenge for environmental and biological disciplines and little is known about structure or potential degradative enzymatic pathways of microbial communities capable of PU biodegradation. Metagenomics provides access to the structure and genetic potential of microbial communities, helping to understand the ecophysiological relationships governing the dynamics of their populations in the environment. Recently, a new approach has been developed that allows the reconstruction of individual genomes of microbial species using physical interactions between sequences within cells [26]. This approach involves Hi-C proximity ligation and yields direct evidence of sequences co-occurrence within a genome, which is used for *de novo* assembly, identification of complete and novel genomes [27] and for testing functional and phylogenetic hypotheses, surpassing other methods for clustering contigs by taxonomic origins [28–30].

To characterize the biodegradation process of the recalcitrant plastic PE-PU by microbial communities, we adopted the commercial water PU dispersion PolyLack^®^ (Sayer Lack, México) that contains a proprietary aromatic polyether-polyurethane-acrylate (PE-PU-A) copolymer and the xenobiotic additives N-methyl 2-pyrrolidone (NMP), isopropanol (IP) 2-butoxyethanol (2-BE), dipropyleneglycol butyl ether (DPGB), and dipropyleneglycol methyl ether (DPGM). In this work, we provide comprehensive chemical and physical evidences for the capacity of a selected landfill microbial community to degrade an aromatic PE-PU-A copolymer and the aforementioned xenobiotic additives, and analyze its structure and phenotypic potential by applying the Hi-C proximity ligation technology. Based on these analyses, we identified a novel microbial landscape that can deal with PE-PU-A and xenobiotics additives degradation and proposed the putative metabolic pathways and genes that can account for these capabilities. This is one of the few studies that combine physical and chemical analyses with metagenomics to elucidate possible metabolic pathways involved in xenobiotics biodegradation, and the first metagenomic analysis of a polyurethane-degrading enriched landfill community. Understanding these pathways will help to design environmental biotechnological strategies that contribute to mitigate plastics and xenobiotics pollution and to achieve a better environmental quality.

## MATERIALS AND METHODS

### Microbiological techniques

The BP8 community, studied in this work, was selected by inoculating deteriorated pieces of PU foam collected at El Bordo Poniente municipal landfill, as previously described [21], into a minimal medium (MM) [10] containing PolyLack^®^ (0.3% v/v), as the sole carbon source (MM-PolyLack). PolyLack^®^ (Sayer Lack, Prod. Num. UB-0810, México) contains a proprietary aromatic PE-PU-A copolymer (≤30% w/v), and the additives NMP (≤6% v/v), 2-BE (≤5% v/v), IP (≤3% v/v), DPGB (≤2% v/v), DPGM (≤1% v/v), and silica (≤3% w/v) [31]. BP8 growth was quantified by dry weight. For that, flasks with MM-PolyLack (25 ml) were inoculated with fresh cells (3 mg/ml) harvested from pre-cultures grown in MM-PolyLack for 48 h at 37°C, 220 rpm. At different incubation times, cells of one flask were harvested, washed three times with phosphate buffer (50 mM, pH 7) and dried to constant weight. Emulsification index (EI24) and cell surface hydrophobicity (CSH) were determined as described [32]. To observe cell-copolymer interactions, cells were fixed with 3% (v/v) glutaraldehyde in phosphate buffer (100 mM, pH 7.4), at 4°C overnight, washed three times, dehydrated with serial dilutions of ethanol, coated with gold and analyzed in a JEOL JSM-5900-LV electron microscope.

### Analytical techniques

Nuclear magnetic resonance spectra from dried PolyLack^®^ dissolved in C5D5N (30 mg/ml) were recorded at 298 K in a Bruker Avance 400 NMR (Billerica, MA, USA) at 400 MHz (^1^H). For most of the analytical techniques, cell-free supernatants (CFS) were obtained by centrifugation at 17 211 × *g* for 10 min, filtered through Whatman grade 41 paper, and dried at 37°C for 5 days. Carbon content was determined in a Perkin Elmer Elemental Analyzer (2400 CHN/O, Series II, Shelton, CT., USA). For gas chromatography coupled to mass spectrometry (GC-MS) analysis, 25 ml CFS were extracted in 6 ml LC-18 cartridges (Supelco) at a flow rate of 2 ml/min, eluted with 2 ml chloroform:methanol (1:1, v/v) and concentrated to 0.5 ml. Samples were injected in an Agilent GC system (7890B, Santa Clara, CA, USA) using two 5%-phenyl-methylpolysiloxane columns (15 m × 250 μm × 0.25 μm). Oven was heated from 50°C to 300°C at 20°C/min, Helium was used as carrier gas at a flow rate of 1 ml/min. The injector temperature was 300°C. For the quantification of additives, pure compounds (Sigma-Aldrich Chemicals ≥98% purity) were used for standard curves. Identification of biodegradation products was performed in an Agilent Quadrupole Mass Analyzer (5977A MSD, Santa Clara, CA, USA) with electronic ionization energy of 1459 EMV and the mass range scanned at 30-550 amu. Scan rate was 2.8 spec/s. Data acquisition was performed with the Enhanced MassHunter software system. Compounds were identified based on mass spectra compared to the NIST database (2002 Library). Fourier transform infrared spectroscopy (FTIR) analyses were performed in a Perkin Elmer spectrometer (Spectrum 400, Waltham, MA, USA) in attenuated total reflection mode; 64 scans with a resolution of 4 cm^−1^ were averaged in the range of 500-4000 cm^−1^, processed and analyzed (Spectrum v6.3.5.0176 software). Derivative thermogravimetric analyses (DTG) were performed in a Perkin Elmer Thermogravimetric Analyzer (TGA 4000, Waltham, MA, USA) on 2.5 mg of dried CFS samples heated 30-500°C at a rate of 20°C/min, under a N2 atmosphere. Differential Scanning Calorimetry (DSC) was performed analyzing 10 mg of dry CFS in a Q2000 (TA Instrument, New Castle, DE, USA) at a rate of 10°C/min, under a nitrogen flow of 50 ml/min, at a 20-600°C range. Gel Permeation Chromatography was performed in a Waters 2695 Alliance Separation Module GPC (Milford, MA, USA) at 30°C in tetrahydrofuran, using a universal column and a flow rate of 0.3 ml/min in CFS. All the analyses were performed at least in three replicates. Controls were non-inoculated MM-PolyLack supernatants similarly processed.

### HI-C proximity ligation based metagenomic analysis

BP8 community cells cultured for 5 days in 50 ml of MM-PolyLack were harvested and washed three times with phosphate buffer. Cells were resuspended in 20 ml TBS buffer with 1% (v/v) formaldehyde (J.T. Baker) (crosslinker) and incubated 30 min with periodic mixing. The crosslinker was quenched with glycine (0.2 g) (Bio-Rad) for 20 min, thereafter cells were centrifuged, lyophilized and frozen at −20°C. For DNA extraction, cell pellets (100 μl) were resuspended in 500 μl of TBS buffer containing 1% (v/v) Triton-X 100 and protease inhibitors [27]. DNA was digested with *Sau*3AI and *Mlu*CI and biotinylated with DNA Polymerase I Klenow fragment (New England Biolabs) followed by ligation reactions incubated for 4 h and then overnight at 70°C to reverse crosslinking. The Hi-C DNA library was constructed by using the HyperPrep Kit (KAPA Biosystems, Wilmington, MA, USA). A shotgun library was also prepared from DNA extracted from non-crosslinked cells using Nextera DNA Sample Preparation Kit (Illumina). The two libraries were paired-end sequenced using NextSeq 500 Illumina platform (Illumina, San Diego, CA, USA). *De novo* metagenome draft assemblies from the raw reads were made using the metaSPAdes assembler [33]. Hi-C reads were then aligned to the contigs obtained from the shotgun library using the Burrows-Wheeler Alignment tool [34] requiring exact read matching. The ProxiMeta algorithm was used to cluster the contigs of the draft metagenome assembly into individual genomes [27]. Additionally, we performed a community taxonomic profiling from shotgun reads using MetaPhlAn tool [35]. Genome completeness, contamination, and other genomic characteristics were evaluated using CheckM pipeline [36]. Phylogenetic analysis was performed using the single copy molecular markers, DNA gyrase subunit A and ribosomal proteins L3 and S5, selected from each deconvoluted genome and compared to homologous sequences from GenBank. Alignments were cured with Gblocks tool (http://phylogeny.lirmm.fr/phylo_cgi/one_task.cgi?task_type=gblocks) and WAG plus G evolutionary models were selected using Smart Model Selection tool [37]. Finally, phylogeny was inferred with the graphical interface of SeaView [38] using the Maximum Likelihood method. To compare genetic relatedness, Average Nucleotide Identity (ANI) between the genomes and the closest phylogenetic neighbors was calculated [39]. Open reading frames were identified using MetaGeneMark [40]. KO assignments (KEGG Orthology) and KEGG pathways reconstruction were performed with GhostKOALA server and KEGG Mapper tool, respectively [41]. All the xenobiotic degradation pathways were manually curated to only report those pathways in which most of the enzymes were encoded in the BP8 metagenome.

### Data availability

Genomes described in this manuscript were deposited to GenBank under Bioproject Accession number: PRJNA488119.

## RESULTS

### Growth and interactions of BP8 cells with PolyLack^®^

The BP8 community cultivated in MM-PolyLack for 25 days exhibited a biphasic growth with a first phase, from 0-13 days, presenting a growth rate (2-4 days) of 0.008 h^−1^ and a second phase, from 13-25 days, with a growth rate (13-20 days) of 0.005 h^−1^. Biomass increased from 0.32 to 2.9 mg/ml and consumed 50.3% of the carbon from the medium at 25 days (Figure 1a). EI24 initial value was 70%, it decreased to 24% at 20 days and increased again to 70%. CSH started at 62% and decreased to 25% at the first growth phase, thereafter it increased to 42% and remained constant until 20 days to increase to 67% at the end of the second phase (Figure 1b). SEM analysis at 10 days of cultivation revealed multiple-sized (0.5-1.5 μm) rod-shaped cells aggregated and attached to copolymer particles (Figure 1c). The changes in CSH and EI24, reflect the complex cell-substrate interactions involved in promoting substrate bioaccessibility and mineralization, as has been observed in bacteria degrading other xenobiotics [42, 43].

**Figure 1.**
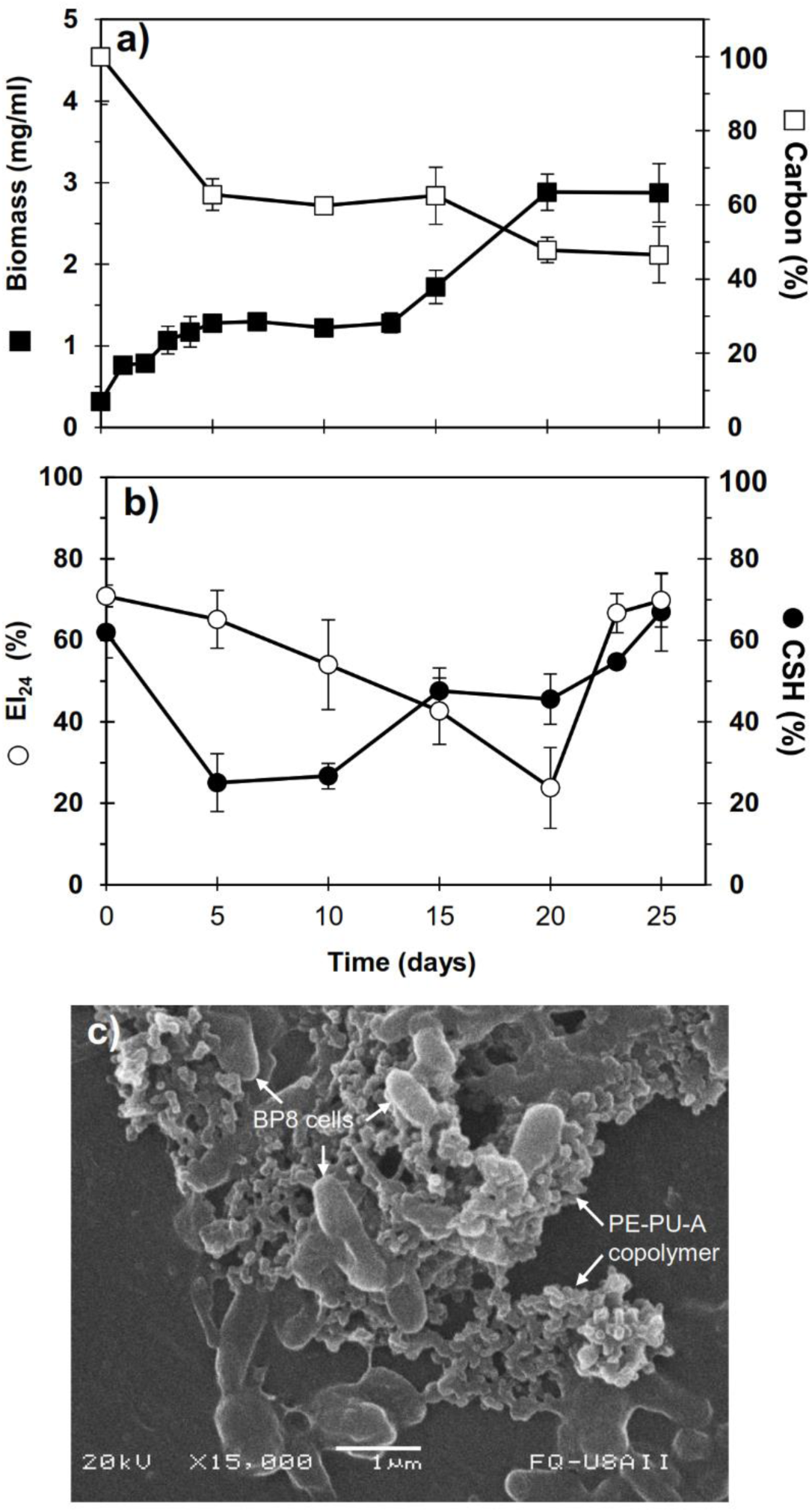
Characteristics of the BP8 community growing in MM-PolyLack. **a)** Growth and carbon consumption, **b)** emulsification index (EI24) and cell surface hydrophobicity (CSH) at different cultivation times; **c)** SEM micrograph of BP8 cells attached to the PE-PU-A copolymer at 10 days of cultivation. Bars represent standard deviation. n=3.

### Chemical and physical changes in PolyLack^®^ components generated by the BP8 community

To characterize the biodegradative activity of the BP8 community on the PolyLack^®^ components, we performed different analytical techniques. GC-MS analysis of the CFS revealed that BP8 metabolized the xenobiotic additives, NMP and IP at the first day of cultivation, 2-BE at the fourth day and DPGM and DPGB were metabolized 85 and 73% respectively at the first day, and remained constant until the end of the experiment (Figure 2). Since the PE-PU-A copolymer structure is unknown, we proposed a hypothetical structure (Figure S1), based on ^1^H-NMR, the manufacturer’s technical sheet and in the most frequently used chemicals for the synthesis of this copolymer [44–46]. Since the first day of cultivation, complex and diverse chemical compounds such as aromatics, nitrogen-containing, ethers, esters, aliphatics, alcohols and organic acids, derived from the copolymer breakdown were observed. During the first 3 days (log phase) the degradation products were low abundant, at 10 days (intermediate lag phase) accumulation occurred, and during the second log phase their abundance decreased. Notably, isocyanates (2,4-toluene diisocyanate (TDI) and methylene diphenyl diisocyanate (MDI)) derivatives were aromatic amines observed maximal at the beginning and diminished throughout the cultivation period (Figure 2, S2), suggesting that metabolization of the urethane groups is being achieved. FTIR of CFS revealed changes in PE-PU-A functional groups. The signal intensity of the C=O stretch from urethane and acrylate carbonyl groups (1 730 cm^−1^) increased at 5 days and lately decreased, suggesting hydrolysis and subsequent catabolism of urethanes and acrylates. The signal for aromatic groups C=C stretch (1 600 cm^−1^) considerably decreased at 20 days, while the signal for aromatic C-C stretch (1 380 cm^−1^) showed variable intensities at different days, and a new C-C signal for aromatics (1 415 cm^−1^) appeared at 20 days, indicating the cleavage of the aromatic rings. The urethane N-H bending plus C-N stretch signal (1 530 cm^−1^) slightly decreased at 15 days and increased at the end of the cultivation time, whereas urethane C-N stretching band (1 231 cm^−1^) significantly increased, indicating urethane attack. Signals associated with urethane C-O-C stretch (1 086 cm^−1^, 1 049 cm^−1^) and C-O-C symmetric stretch (977 cm^−1^) decreased during the cultivation period, indicating microbial activity on the ether groups. The signal for the acrylate’s vinyl group C=C-H out of plane (850 cm^−1^) decreased at 20 days, indicating the cleavage of the acrylate component. Also, the aliphatic chain signals (704 and 520 cm^−1^) decreased during the cultivation period (Figure 3a). DTG thermograms exhibited four stages of thermal decomposition corresponding to the functional groups of the copolymer. Stages II and IV, for urethane and ether groups respectively, reduced their masses at early cultivation times, while stage III, for esters, steadily kept reducing its mass during the whole experimental period. Interestingly, stage I, which accounts for low molecular weight compounds, in this case biodegradation products, showed a fluctuating behavior that increased at 10 days, and decreased afterwards (Figure 3b). DSC analysis of the copolymer showed multiple thermal transitions revealing complex microstructures: the glass transition temperature (Tg: 50.2°C) reflects the proportion of soft and hard segments; the three melting temperatures (Tm-I: 70°C, Tm-II: 210.6°C, Tm-III: 398.1°C) are associated with the hard segments of the polymer and the crystallization temperature (Tc: 459.6°C) is attributed to heat-directed crystallization of copolymer chains [47, 48] (Figure 3c). BP8 biodegradative activity caused Tg decrease (46.2°C), changes in Tms, and strong decrease in Tc area, indicating that BP8 disrupts both, the soft and the hard segments (associated with urethane groups) (Figure 3, Table S1). GPC analysis showed that the number-average molecular weight of the copolymer decreased 35.6% and the PDI increased to values higher than 2, at 25 days of cultivation with BP8 (Table S2). All these results indicate that the degradative activity of the BP8 community generates changes in the soft and hard segments of the copolymer microstructure resulting from the attack to the different functional groups, including the more recalcitrant ether and urethane groups.

**Figure 2.**
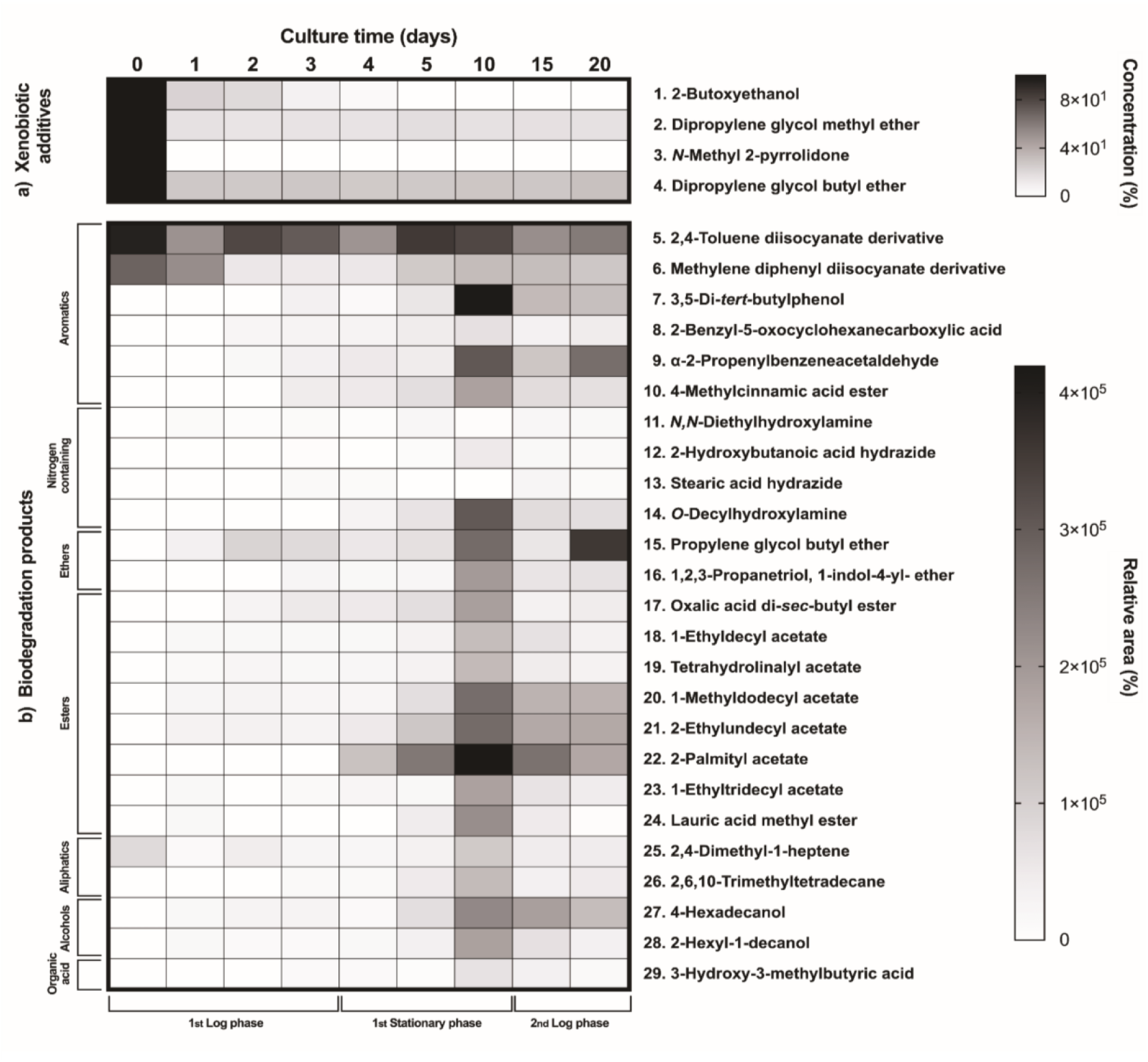
Xenobiotic additives consumed (a) and PE-PU-A biodegradation products generated (b) by the BP8 community. Cell-free supernatants were extracted at differents culture times with chloroform:methanol and analyzed by GC-MS. **a)** Additives were quantified using standard curves for each compound and **b)** biodegradation products by analyzing their relative areas in independent chromatograms. n=3. Compounds with mass spectra similarity values over 700 were considered the same compounds of the Library hits. The numbers in the compounds correspond to signals in the chromatograms of Figure S2.

**Figure 3.**
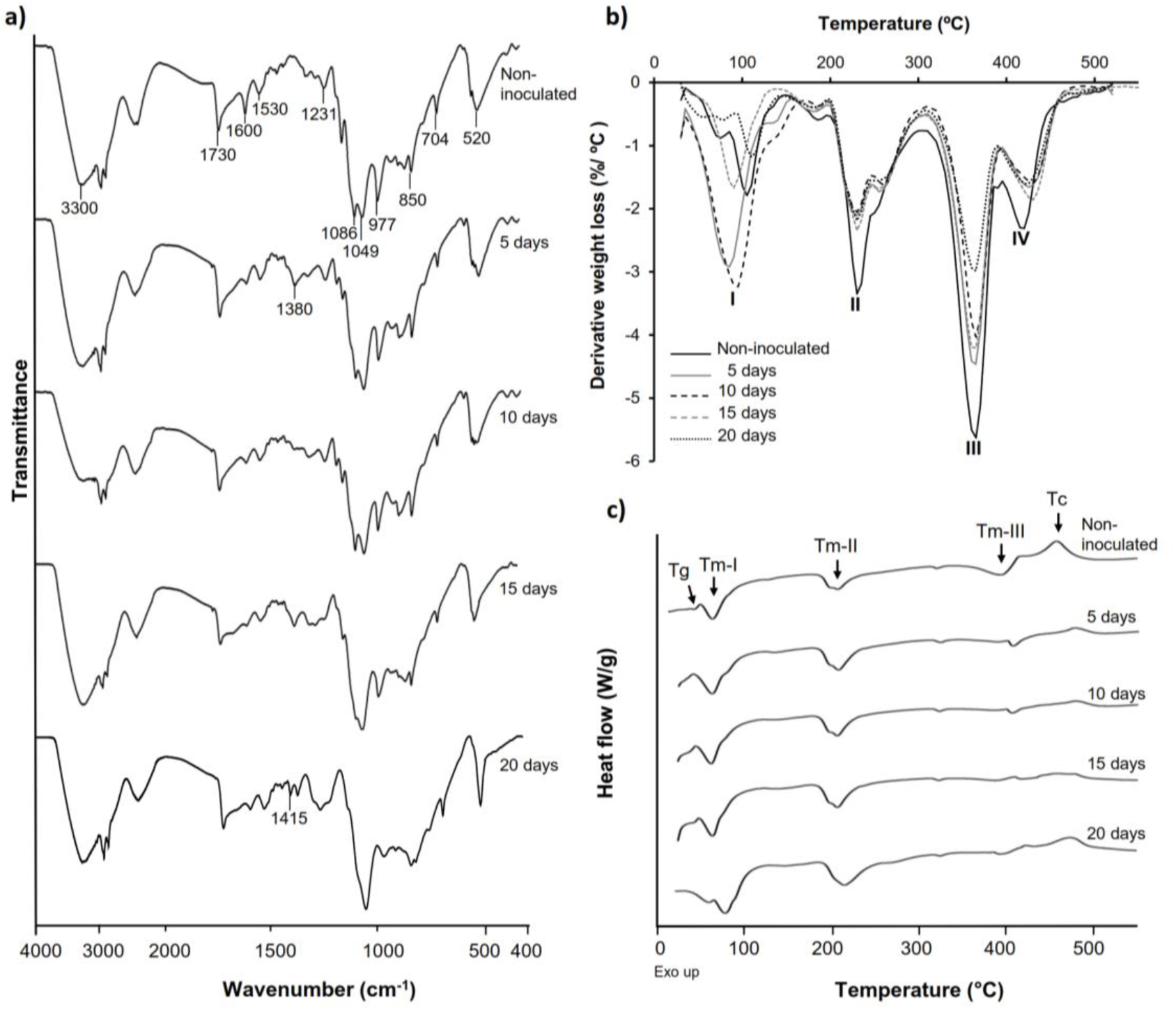
Physical and chemical analyses of the aromatic PE-PU-A copolymer after incubation with the BP8 community. **a)** FTIR spectra. **b)** DTG analysis. Thermal degradation stages correspond to the following functional groups: I. Low molecular weight compounds, II. Urethane, III. Ester, IV. Ether; **c)** DSC analysis. Glass transition temperature (Tg) represents the relative amount of soft and hard segments; melting temperatures, Tm-I, Tm-II and Tm-III are associated with hard domains, and crystallization temperature (Tc) represents heat-directed crystallization of copolymer chains.

### Community structure and metagenomic deconvolution of the BP8 community

Analysis of the BP8 community taxonomic profile with MetaPhlAn, by using 17 282 414 reads, detected five bacterial orders (abundance), *Rhodobacterales* (83%), *Rhizobiales* (8.9%), *Burkholderiales* (6.8%), *Actinomycetales* (0.83%), *Sphingobacteriales* (0.08%), and one viral order *Caudovirales* (0.33%). Bacteria included 16 genera, being the most abundant *Paracoccus* (83.9%) and *Ochrobactrum* (8.7%) (Figure S3). *De novo* assembly of the shotgun metagenome sequences generated 3 072 contigs with a total length of 17 618 521 bp. Alignment of Hi-C reads to this assembly allowed the deconvolution of five genome clusters, three near complete drafts (>95%), and two substantially complete drafts (89 and 71%) [36] (Table S3). The phylogenetic analysis showed well-supported clades within *Paracoccus*, *Chryseobacterium*, *Parapedobacter*, a member of the *Microbacteriaceae* family, and *Ochrobactrum intermedium* (Figure S4). The deconvoluted genomes of *Paracoccus* sp. BP8 and *O. intermedium* BP8.5 showed low novelty scores and high ANI values compared to their closest phylogenetic relatives, while *Chryseobacterium* sp. BP8.2, *Parapedobacter* sp. BP8.3 and the *Microbacteriaceae* bacterium BP8.4 showed high novelty scores and low ANI values (<95%) indicating they are new species. GC content and genomes’ sizes were similar to the closest relatives except for the *O. intermedium* BP8.5 genome size, probably because of the low genome completeness (Table S3, S4).

### Analysis of the xenobiotic metabolism encoded in the BP8 metagenome

In all the genomes, except in *O. intermedium* BP8.5, the genes and proteins assigned were in the range reported for the phylogenetically related members (Table S3, S4). Reconstruction of the metabolic pathways encoded in the BP8 metagenome was performed with 18 386 ORFs from which 8 637 were annotated into KEGG Orthology groups and the rest was not assigned to any orthologous functional category. Analysis of the BP8 xenobiotic metabolism identified 215 sequences encoding 59 unique proteins participating in pathways for benzoate (ko00362), fluorobenzoate (ko00364), aminobenzoate (ko00627), chlorocyclohexane and chlorobenzene (ko00361), and n-alkanes (ko00071) degradation. The most relevant enzymes are listed in Table 1. The genes for benzoate metabolism include all the enzymes for benzoate and 4-methoxybenzoate activation as well as 4-methoxybenzoate monooxygenase, a O-demethylating enzyme that transforms methoxybenzoate to hydroxybenzoate, and for their subsequent transformation to β-ketoadipate (first 18 EC numbers in Table 1). Two genes encoding carboxymethylene butanolidase that cleavages the ring of cyclic ester dienelactone to produce maleylacetate, acting on the fluorobenzoate and chlorocyclohexane and chlorobenzene metabolisms, were identified. Genes encoding enzymes for the aminobenzoate pathway, such as 4-hydroxybenzoate decarboxylase that participates in the transformation of phenol into hydroxybenzoate, amidase that transforms benzamide into benzoate, and benzoyl phosphate phospohydrolase that converts benzoyl phosphate into benzoate, were identified. All the genes encoding enzymes needed for chlorocyclohexane and chlorobenzene degradation, the specific 2,4-dichlorophenol 6-monooxygenase, the enzymes that transform 4-chlorophenol to cis-acetylacrylate (EC1.13.11.1, EC5.5.1.1 and EC3.1.1.45), and the 2-haloacid dehalogenase, which eliminates halogens from alkanes, were found. Likewise, genes encoding enzymes for n-alkanes degradation (Table 1 Alkanes metabolism), as well as all the enzymes for beta-oxidation were also detected.

**Table 1.**
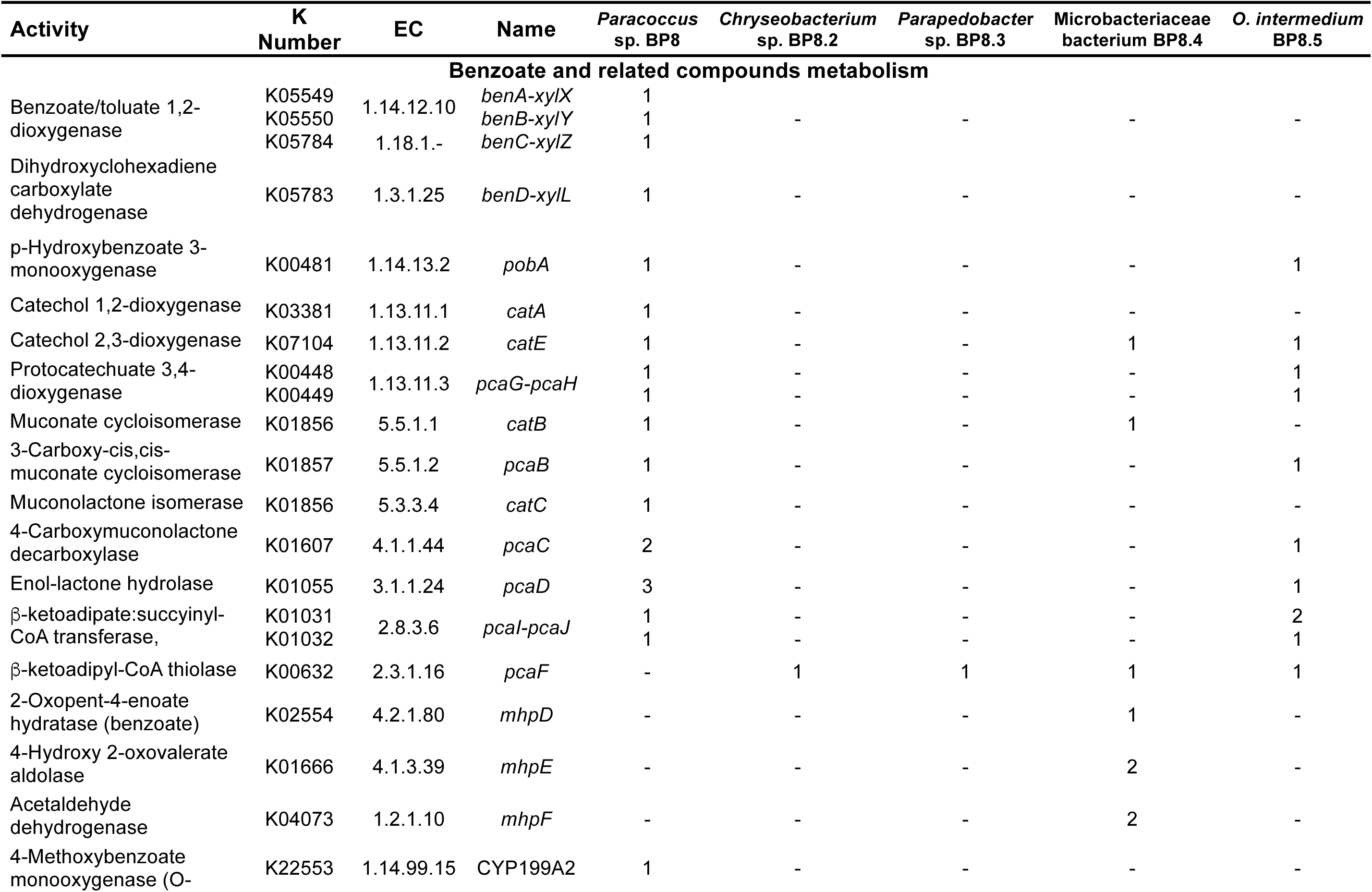
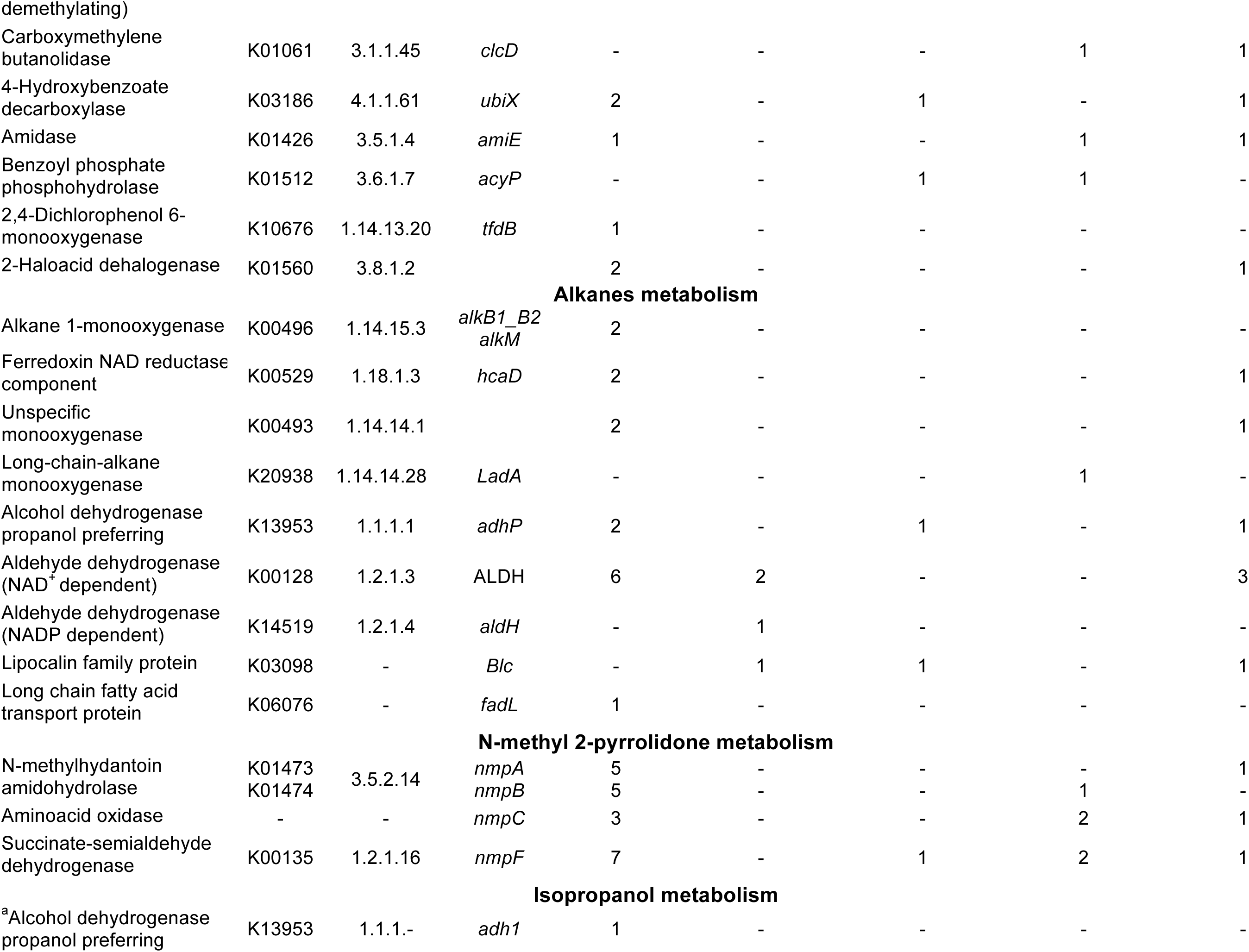
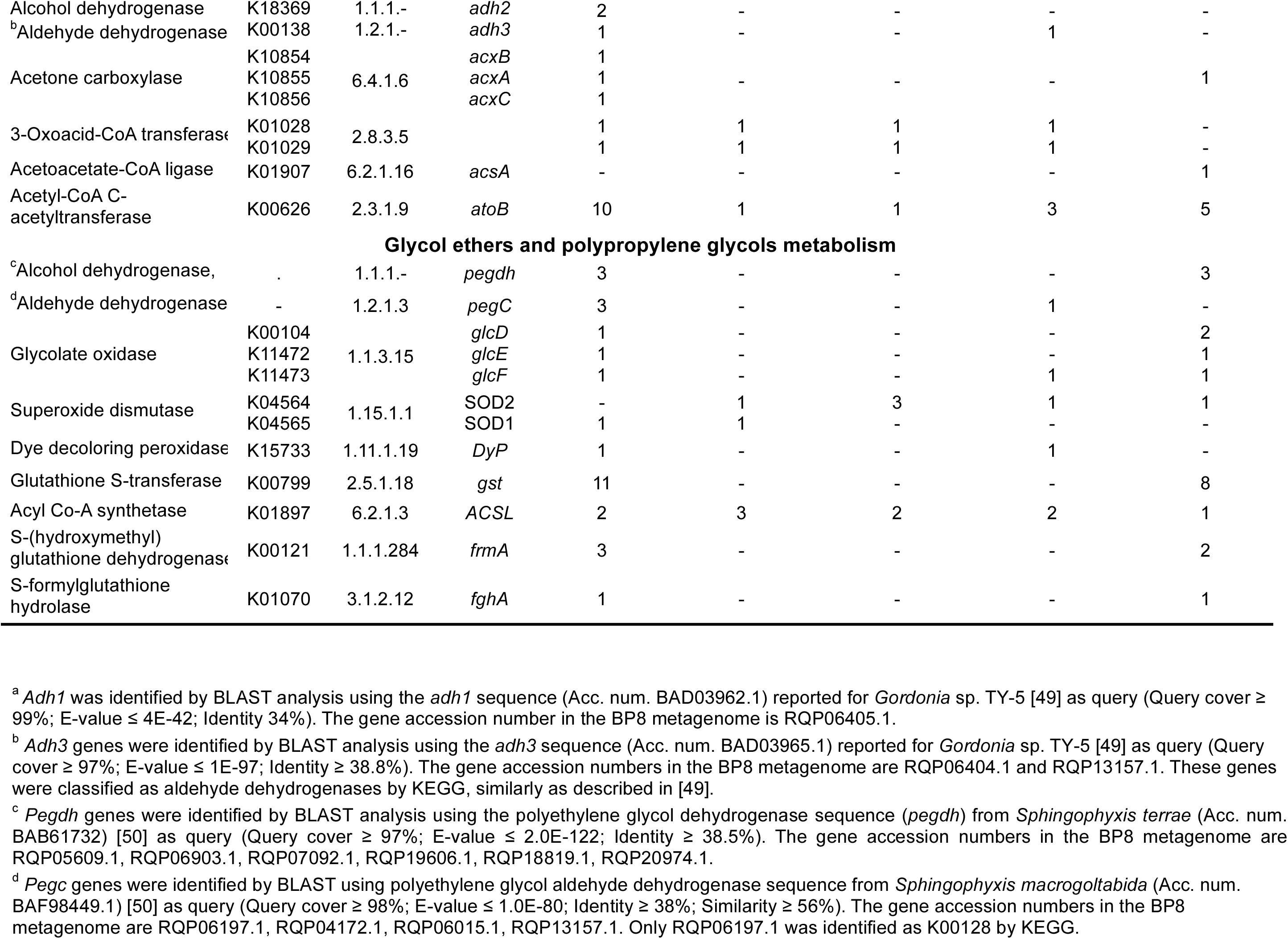
Distribution of genes encoding relevant proteins involved in xenobiotics degradation in the BP8 metagenome.

| Activity | K Number | EC | Name | *Paracoccus*<br>sp. BP8 | *Chryseobacterium*<br>sp. BP8.2 | *Parapedobacter*<br>sp. BP8.3 | *Microbacteriaceae*<br>bacterium BP8.4 | *O. intermedium*<br>BP8.5 |
| --- | --- | --- | --- | --- | --- | --- | --- | --- |
| **Benzoate and related compounds metabolism** |  |  |  |  |  |  |  |  |
| Benzoate/toluate 1,2-dioxygenase | K05549 | 1.14.12.10 | *benA-xylX* | 1 | - | - | - | - |
|  | K05550 |  | *benB-xylY* | 1 |  |  |  |  |
|  | K05784 | 1.18.1.- | *benC-xylZ* | 1 |  |  |  |  |
| Dihydroxycyclohexadiene carboxylate dehydrogenase | K05783 | 1.3.1.25 | *benD-xylL* | 1 | - | - | - | - |
| p-Hydroxybenzoate 3-monooxygenase | K00481 | 1.14.13.2 | *pobA* | 1 | - | - | - | 1 |
| Catechol 1,2-dioxygenase | K03381 | 1.13.11.1 | *catA* | 1 | - | - | - | - |
| Catechol 2,3-dioxygenase | K07104 | 1.13.11.2 | *catE* | 1 | - | - | 1 | 1 |
| Protocatechuate 3,4-dioxygenase | K00448 | 1.13.11.3 | *pcaG-pcaH* | 1 | - | - | - | 1 |
|  | K00449 |  |  | 1 | - | - | - | 1 |
| Muconate cycloisomerase | K01856 | 5.5.1.1 | *catB* | 1 | - | - | 1 | - |
| 3-Carboxy-cis,cis-muconate cycloisomerase | K01857 | 5.5.1.2 | *pcaB* | 1 | - | - | - | 1 |
| Muconolactone isomerase | K01856 | 5.3.3.4 | *catC* | 1 | - | - | - | - |
| 4-Carboxymuconolactone decarboxylase | K01607 | 4.1.1.44 | *pcaC* | 2 | - | - | - | 1 |
| Enol-lactone hydrolase | K01055 | 3.1.1.24 | *pcaD* | 3 | - | - | - | 1 |
| β-keto adipate:succinyl-CoA transferase, | K01031 | 2.8.3.6 | *pcaI-pcaJ* | 1 | - | - | - | 2 |
|  | K01032 |  |  | 1 | - | - | - | 1 |
| β-keto adipyl-CoA thiolase | K00632 | 2.3.1.16 | *pcaF* | - | 1 | 1 | 1 | 1 |
| 2-Oxopent-4-enoate hydratase (benzoate) | K02554 | 4.2.1.80 | *mhpD* | - | - | - | 1 | - |
| 4-Hydroxy 2-oxovalerate aldolase | K01666 | 4.1.3.39 | *mhpE* | - | - | - | 2 | - |
| Acetaldehyde dehydrogenase | K04073 | 1.2.1.10 | *mhpF* | - | - | - | 2 | - |
| 4-Methoxybenzoate monooxygenase (O- | K22553 | 1.14.99.15 | CYP199A2 | 1 | - | - | - | - |
| Carboxymethylene butanolidase | K01061 | 3.1.1.45 | *clcD* | - | - | - | 1 | 1 |
| 4-Hydroxybenzoate decarboxylase | K03186 | 4.1.1.61 | *ubiX* | 2 | - | 1 | - | 1 |
| Amidase | K01426 | 3.5.1.4 | *amiE* | 1 | - | - | 1 | 1 |
| Benzoyl phosphate phosphohydrolase | K01512 | 3.6.1.7 | *acyP* | - | - | 1 | 1 | - |
| 2,4-Dichlorophenol 6-monooxygenase | K10676 | 1.14.13.20 | *tfdB* | 1 | - | - | - | - |
| 2-Haloacid dehalogenase | K01560 | 3.8.1.2 |  | 2 | - | - | - | 1 |
| Alkanes metabolism |  |  |  |  |  |  |  |  |
| Alkane 1-monooxygenase | K00496 | 1.14.15.3 | *alkB1_B2*<br>*alkM* | 2 | - | - | - | - |
| Ferredoxin NAD reductase component | K00529 | 1.18.1.3 | *hcaD* | 2 | - | - | - | 1 |
| Unspecific monooxygenase | K00493 | 1.14.14.1 |  | 2 | - | - | - | 1 |
| Long-chain-alkane monooxygenase | K20938 | 1.14.14.28 | *LadA* | - | - | - | 1 | - |
| Alcohol dehydrogenase propanol preferring | K13953 | 1.1.1.1 | *adhP* | 2 | - | 1 | - | 1 |
| Aldehyde dehydrogenase (NAD<sup>+</sup> dependent) | K00128 | 1.2.1.3 | ALDH | 6 | 2 | - | - | 3 |
| Aldehyde dehydrogenase (NADP dependent) | K14519 | 1.2.1.4 | *aldH* | - | 1 | - | - | - |
| Lipocalin family protein | K03098 | - | *Blc* | - | 1 | 1 | - | 1 |
| Long chain fatty acid transport protein | K06076 | - | *fadL* | 1 | - | - | - | - |
| N-methyl 2-pyrrolidone metabolism |  |  |  |  |  |  |  |  |
| N-methylhydantoin amidohydrolase | K01473 | 3.5.2.14 | *nmpA* | 5 | - | - | - | 1 |
|  | K01474 |  | *nmpB* | 5 | - | - | 1 | - |
| Aminoacid oxidase | - | - | *nmpC* | 3 | - | - | 2 | 1 |
| Succinate-semialdehyde dehydrogenase | K00135 | 1.2.1.16 | *nmpF* | 7 | - | 1 | 2 | 1 |
| Isopropanol metabolism |  |  |  |  |  |  |  |  |
| <sup>a</sup>Alcohol dehydrogenase propanol preferring | K13953 | 1.1.1.- | *adh1* | 1 | - | - | - | - |
| Alcohol dehydrogenase | K18369 | 1.1.1.- | *adh2* | 2 | - | - | - | - |
| <sup>b</sup>Aldehyde dehydrogenase | K00138 | 1.2.1.- | *adh3* | 1 | - | - | 1 | - |
|  | K10854 |  | *acxB* | 1 |  |  |  |  |
| Acetone carboxylase | K10855 | 6.4.1.6 | *acxA* | 1 | - | - | - | 1 |
|  | K10856 |  | *acxC* | 1 |  |  |  |  |
| 3-Oxoacid-CoA transferase | K01028 |  |  | 1 | 1 | 1 | 1 | - |
|  | K01029 | 2.8.3.5 |  | 1 | 1 | 1 | 1 | - |
| Acetoacetate-CoA ligase | K01907 | 6.2.1.16 | *acsA* | - | - | - | - | 1 |
| Acetyl-CoA C-acetyltransferase | K00626 | 2.3.1.9 | *atoB* | 10 | 1 | 1 | 3 | 5 |
| **Glycol ethers and polypropylene glycols metabolism** |  |  |  |  |  |  |  |  |
| <sup>c</sup>Alcohol dehydrogenase, | . | 1.1.1.- | *pegdh* | 3 | - | - | - | 3 |
| <sup>d</sup>Aldehyde dehydrogenase | - | 1.2.1.3 | *pegC* | 3 | - | - | 1 | - |
|  | K00104 |  | *glcD* | 1 | - | - | - | 2 |
| Glycolate oxidase | K11472 | 1.1.3.15 | *glcE* | 1 | - | - | - | 1 |
|  | K11473 |  | *glcF* | 1 | - | - | 1 | 1 |
| Superoxide dismutase | K04564 | 1.15.1.1 | SOD2 | - | 1 | 3 | 1 | 1 |
|  | K04565 |  | SOD1 | 1 | 1 | - | - | - |
| Dye decoloring peroxidase | K15733 | 1.11.1.19 | *DyP* | 1 | - | - | 1 | - |
| Glutathione S-transferase | K00799 | 2.5.1.18 | *gst* | 11 | - | - | - | 8 |
| Acyl Co-A synthetase | K01897 | 6.2.1.3 | *ACSL* | 2 | 3 | 2 | 2 | 1 |
| S-(hydroxymethyl) glutathione dehydrogenase | K00121 | 1.1.1.284 | *frmA* | 3 | - | - | - | 2 |
| S-formylglutathione hydrolase | K01070 | 3.1.2.12 | *fghA* | 1 | - | - | - | 1 |

<sup>a</sup> *Adh1* was identified by BLAST analysis using the *adh1* sequence (Acc. num. BAD03962.1) reported for *Gordonia* sp. TY-5 [49] as query (Query cover ≥ 99%; E-value ≤ 4E-42; Identity 34%). The gene accession number in the BP8 metagenome is RQP06405.1.
<sup>b</sup> *Adh3* genes were identified by BLAST analysis using the *adh3* sequence (Acc. num. BAD03965.1) reported for *Gordonia* sp. TY-5 [49] as query (Query cover ≥ 97%; E-value ≤ 1E-97; Identity ≥ 38.8%). The gene accession numbers in the BP8 metagenome are RQP06404.1 and RQP13157.1. These genes were classified as aldehyde dehydrogenases by KEGG, similarly as described in [49].
<sup>c</sup> *Pegdh* genes were identified by BLAST analysis using the polyethylene glycol dehydrogenase sequence (*pegdh*) from *Sphingophyxis terrae* (Acc. num. BAB61732) [50] as query (Query cover ≥ 97%; E-value ≤ 2.0E-122; Identity ≥ 38.5%). The gene accession numbers in the BP8 metagenome are RQP05609.1, RQP06903.1, RQP07092.1, RQP19606.1, RQP18819.1, RQP20974.1.
<sup>d</sup> *Pegc* genes were identified by BLAST using polyethylene glycol aldehyde dehydrogenase sequence from *Sphingophyxis macrogoltabida* (Acc. num. BAF98449.1) [50] as query (Query cover ≥ 98%; E-value ≤ 1.0E-80; Identity ≥ 38%; Similarity ≥ 56%). The gene accession numbers in the BP8 metagenome are RQP06197.1, RQP04172.1, RQP06015.1, RQP13157.1. Only RQP06197.1 was identified as K00128 by KEGG.

### BP8 community phenotypic potential to biodegrade the xenobiotic additives present in PolyLack^®^

*NMP degradation.* Genes encoding putative proteins for NMP degradation, with significant similarity (>40%) to the enzymes of *Alicycliphilus denitrificans* BQ1 [52] were identified in several BP8 genomes (Table 1). However, only in *Paracoccus* sp. BP8 a gene cluster (RQP05666.1-RQP05671.1) comparable to the BQ1 *nmp* cluster was identified. *Isopropanol degradation.* Genes encoding proteins with significant similarity to NAD^+^-dependent secondary ADH with capability to oxidize IP to acetone were identified in the BP8 metagenome [49], but not the genes encoding the enzymes for the oxidative transformation of acetone. However, the three genes encoding acetone carboxylase, that transforms acetone into acetoacetate, were identified, as well as the enzymes that convert acetoacetate into acetoacetyl-CoA and this to acetyl-CoA are also encoded in the BP8 metagenome (Figure 4a, Table 1). *Glycol ethers degradation.* In the BP8 metagenome, homologous genes to PEG-degrading ADHs and ALDHs [50, 51], and diverse enzymes that could attack the ether bonds, such as glycolate oxidase (RQP04511.1, RQP04512.1, RQP04513.1, RQP11464.1, RQP19624.1, RQP19625.1, RQP16322.1, RQP16256.1), dye decoloring peroxidase (RQP04907.1, RQP09154.1) and superoxide dismutase (RQP04715.1, RQP13424.1, RQP09887.1, RQP11889.1, RQP18047.1, RQP18034.1, RQP09190.1, RQP20377.1), as well as genes encoding enzymes involved in glutathione metabolism, which have been proposed to participate in PEG metabolism [53] were identified (Figure 4b, Table 1).

**Figure 4.**
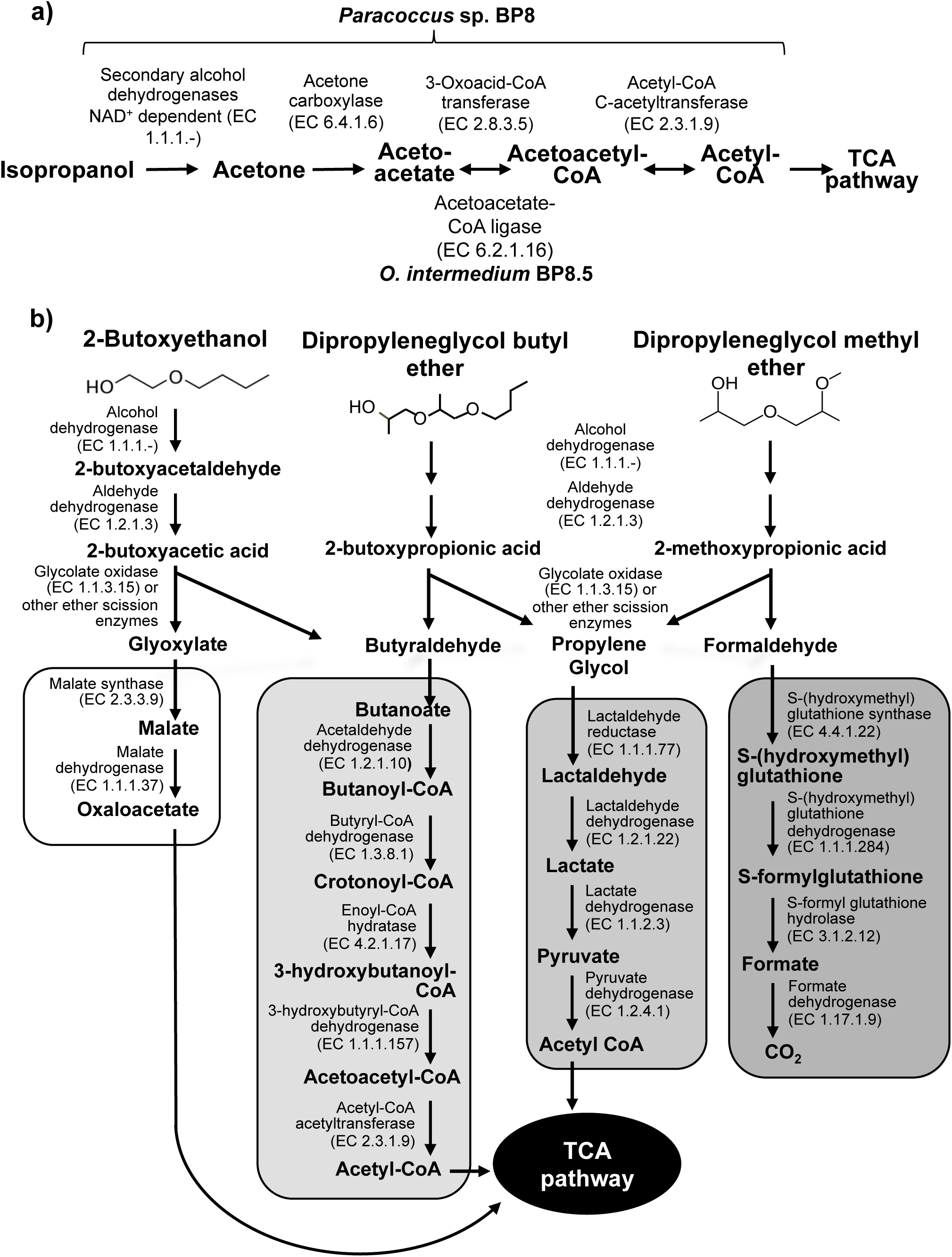
Potential degradation pathways for isopropanol (a) and glycol ethers (b) encoded in the BP8 metagenome. **a)** *Paracoccus* sp. BP8 genome encodes ADH enzymes that can oxidize IP to acetone, but no genes encoding enzymes for acetone metabolism were found. Instead, the genes encoding the three subunits of acetone carboxylase that reductively transforms acetone to acetoacetate were found. Acetoacetate can be transformed to acetoacetyl-CoA by 3-oxoacid-CoA transferase activity, present in *Paracoccus* sp. BP8 or by acetoacetate-CoA ligase, present in O. intermedium BP8.5. Acetoacetyl CoA is transformed by acetyl-CoA C-acetyltransferase to acetyl CoA, that enters the TCA pathway encoded in the BP8 metagenome (See Table 1). **b)** Degradation of 2-BE could be carried out by subsequent oxidations of the hydroxy terminal group by PEG-DH and PEG-ALDH, followed by scission of the ether bond by glycolate oxidase or other ether scission enzymes to produce glyoxylate and butyraldehyde [5, 65]. Glyoxylate would be funneled to the glyoxylate metabolism (white rectangle) and butyraldehyde to the butanoate metabolism (light gray rectangle). DPGB and DPGM can also be degraded by initial oxidation of the hydroxy terminal groups and further be ether-cleaved by ether scission enzymes. The products of these processes would be butyraldehyde an propylene glycol from DPGB and propylene glycol and formaldehyde from DPGM. Propylene glycol can be funneled to the pyruvate metabolism (medium gray rectangle) and formaldehyde can be transformed by the methane metabolism (dark gray rectangle), Genes encoding homologs for PEG-DH and PEG-ALDH (pegdh and pegc) from *Sphyngophyxis terrae* and S. macrogoltabida, the three subunits of glycolate oxidase (glcD, glcE, glcF) and other possible ether scission enzymes were identified in *Paracoccus* sp. BP8 (See Table 1). Pathways for glyoxylate, butanoate, pyruvate and methane metabolisms as well as the TCA pathway were fully reconstructed from the BP8 metagenome based on KEGG annotated genes, using KEGG Mapper.

### BP8 community phenotypic potential to degrade polyurethane

Genes encoding PU-esterases verified for PU degradation [54–56] and confirmed carbamate-hydrolyzing enzymes *i.e.* arylamidase A [57], amidase [58], urethanase [59], and carbaryl hydrolase [60], were searched by standalone BLASTP analyses. Six and five sequences with similarity to PU-esterases and carbamate hydrolases were retrieved from the BP8 metagenome, respectively (Table 2). We also identified genes encoding ureases (EC3.5.1.5), suggested to act on PU degradation [61], in *Parapedobacter* sp. BP8 (RQP19536.1, RQP19537.1 RQP19538.1) and *O. intermedium* BP8.5 (RQP17756.1, RQP17448.1, RQP17449.1, RQP17450.1) genomes.

**Table 2.** Esterases and carbamate hydrolyzing enzymes encoded in the BP8 metagenome.

| Enzyme<br>Query Organism<br>(Accession number) | E.C.<br>num. | Amino<br>acids in<br>the query | Hit in the BP8<br>metagenome | E value/<br><sup>a</sup>Identity/<br>Similarity | Amino<br>acids in<br>the hit | Reference |
| --- | --- | --- | --- | --- | --- | --- |
| Polyurethane esterase *Delftia acidovorans* (BAA76305) | 3.1.1.6 | 548 | *Parapedobacter* sp. BP8.3 (RQP17780.1) | 1.0E-07/<br>32%/ 48% | 640 | [54] |
| Polyurethanase esterase A<br>*Pseudomonas chlororaphis*<br>(AAD22743) | 3.1.1.- | 617 | *Paracoccus* sp. BP8 (RQP07762.1) | 4.0E-16/<br>34%/ 50% | 783 | [55] |
|  |  |  | *Paracoccus* sp. BP8 (RQP06646.1) | 2.0E-14/<br>33%/ 46% | 980 |  |
|  |  |  | *Paracoccus* sp. BP8 (RQP04598.1) | 7.0E-12/<br>30%/ 43% | 854 |  |
|  |  |  | *Paracoccus* sp. BP8 (RQP06839.1) | 5.0E-12/<br>32%/ 43% | 612 |  |
| Esterase CE_Ubrb<br>uncultured bacterium<br>(SIP63154) | 3.1.1.- | 295 | *Microbacteriaceae* bacterium BP8.4 (RQP12977.1) | 8.0E-05/<br>35%/ 48% | 309 | [56] |
| Arylamidase A<br>*Paracoccus huijuniae*<br>(AEX92978) | 3.4.11.2 | 465 | *Paracoccus* sp. BP8 (RQP04489.1) | 6.0E-25/<br>41%/ 52% | 471 | [57] |
| Amidase<br>*Ochrobactrum* sp. TCC-2<br>(ANB41810) | 3.5.1.4 | 474 | *Microbacteriaceae* bacterium BP8.4 (RQP11486.1) | 2.0E-47/<br>35%/49% | 475 | [58] |
|  |  |  | *O. intermedium* BP8.5 (RQP19215.1) | 6.0E-24/<br>39%/50% | 326 |  |
| Urethanase<br>*Lysinibacillus fusiformis*<br>(KU353448) | 3.5.1.4 | 472 | *Microbacteriaceae* bacterium BP8.4 (RQP12064.1) | 1.0E-60/<br>32%/ 48% | 499 | [59] |
| Carbaryl hydrolase *cahA*<br>*Arthrobacter* sp. RC100<br>(BAC15598) | 3.6.3.5 | 506 | *Paracoccus* sp. BP8 (RQP06118.1) | 1.0E-15/<br>35%/ 48% | 326 | [60] |

<sup>a</sup>Only genes with identity $\geq 30\%$ are presented.

## DISCUSSION

To elucidate the mechanisms that landfill microbial communities perform to degrade the recalcitrant PE-PU plastic, here we studied the degradative activity of the BP8 microbial community that was selected because of its capability to grow in PolyLack^®^, a WPUD that contains a proprietary PE-PU-A copolymer and several xenobiotic additives (NMP, IP, 2-BE, DPGB and DPGM). Chemical and physical analyses demonstrated that BP8 consumes the additives and breaks the copolymer whereas Hi-C based metagenomic analysis allowed us to unveil the phenotypic potential to degrade PU and xenobiotics of five deconvoluted genomes from the community. The diauxic growth of BP8 observed during 25 days of cultivation in MM-PolyLack suggested that two different metabolic processes were involved in degrading the components of the WPUD. We hypothesized that the additives were consumed during the first phase whereas the copolymer was broken during the second one. However, the biomass increment and the carbon decrease observed in the first growth phase (Figure 1a) resulted not only from additive consumption, but also from the copolymer breakdown (Figures 2, 3, S2, Tables S1, S2). These observations indicate that the diauxic growth is the result of simultaneous degradation of additives and copolymer and that microbial enrichment could have selected a more effective PU-degrading community that accounts for the second exponential growth phase. Further studies to demonstrate this possibility are being undertaken.

Exploring the BP8 metagenome, genes encoding enzymes presumably involved in the degradation of the PolyLack^®^ additives were identified in several of the deconvoluted genomes. Genes for NMP degradation, similar to the ones reported for *A. denitrificans* BQ1 [52] were identified in the *Paracoccus* sp. BP8 genome. *Paracoccus* strains able to utilize NMP as carbon source have been reported [62], but the genes sustaining this capability have not been described. IP biodegradation occurs by oxidative pathways in *P. denitrificans* GH3 and *Gordonia* sp. TY-5. In these strains, IP is transformed by NAD^+^-dependent secondary ADH into acetone that is oxidized by a specific monooxygenase to produce methyl acetate, which is transformed to acetic acid and methanol [49, 63]. However, the enzymes for metabolizing acetone by these reactions are not encoded in the BP8 metagenome. Instead, genes encoding enzymes for acetone carboxylation, to produce acetoacetate (acetone carboxylase), and for its subsequent transformation to acetoacetyl-CoA by 3-oxoacid-CoA transferase and thereafter to acetyl-CoA by acetyl-CoA C-acetyltransferase [64] were identified (Figure 4a, Table 1). The possibility that IP degradation occurs by transformation to acetyl-CoA, via acetone in BP8 is supported by the observation that in the *Paracoccus* sp. BP8 genome, a gene encoding an ADH (RQP05888.1), homologous to the *Gordonia* sp. TY-5 *adh*2, and genes encoding the acetone carboxylase subunits (RQP05866.1, RQP05867.1, RQP05889.1) are contiguously located. Adjacent to these genes, a sequence encoding a sigma-54-dependent transcriptional regulator (RQP05868.1) was observed, suggesting an operon-like organization. This presumptive IP degradative operon has not been described in any other bacteria. Degradation of 2-BE, DPGM and DPGB, the glycol ethers present in PolyLack^®^, has not been reported in bacteria. Degradation pathways for PEG and PPG reported in *Sphingomonads* species and *Microbacterium* (formerly *Corynebacterium*) sp. No. 7 [5, 65, 66–68] show similar reactions where the glycols’ hydroxyl terminal groups are sequentially oxidized by specific ADHs and ALDHs to produce aldehydes, and thereafter carboxylic acids [50, 51], suggesting a widespread strategy for glycol ethers metabolism in prokaryotes. Nevertheless, few enzymes involved in scission of ether bonds, present in these compounds, have been identified in bacteria. A glycolic acid oxidase [69] and a glycolic acid dehydrogenase [70] have been reported acting on PEG, although several other enzymes such as superoxide dismutase, monooxygenase, ether hydrolase, carbon-oxygen lyase, peroxidase and laccase have been suggested [5]. Homolog genes for specific ADHs and ALDHs were identified in the *Paracoccus* sp. BP8 genome (Table 1). Therefore, we hypothesize that 2-BE can be oxidized to 2-butoxyacetic acid, DPGM to 2-methoxypropionic acid, which has been reported as a metabolite in the degradation of DPGM by rats [71], and DPGB to 2-butoxypropionic acid (Figure 4b). In *Paracoccus* sp. BP8, and in other genomes of the BP8 community, genes encoding glycolate oxidase, dye decoloring peroxidase, 4-methoxybenzoate monooxygenase and unspecific monooxygenase, which could account for the ether scission of the aforementioned carboxylic acids, were identified (Table 1). The cleavage of the carboxylates produced by ALDHs would generate the metabolizable intermediaries glyoxylate, butyraldehyde, propylene glycol and formaldehyde (Figure 4b). Glyoxylate can be funneled to the glyoxylate metabolism, butyraldehyde to the butanoate metabolism, propylene glycol to the pyruvate metabolism, by lactaldehyde and lactate dehydrogenases as suggested in *P. yeei* TT13 [72], and formaldehyde can be channeled to the formate metabolism where glutathione-dependent enzymes could oxidize it to formate and thereafter to CO2 (Figure 4b, Table 1). All the enzymes for the aforesaid metabolic pathways are encoded in the BP8 metagenome. Additionally, in PEG metabolism, long chains of PEG-carboxylate can be processed by acyl-CoA synthetase and glutathione-S transferase forming glutathione-conjugates [53]. Although these reactions would not be needed for glycol ethers catabolism, they could be required for the degradation of long polypropylene glycol moieties that are part of the PE-PU-A copolymer (Figure S1).

By using different analytical techniques, we demonstrate that the BP8 community attacks the main functional groups of the PE-PU-A copolymer; from the more enzymatically susceptible ester bonds, present in acrylate and carbamate, to the more recalcitrant C-C from aliphatics and aromatics, C-N from urethane, and C-O-C from ether bonds of polypropylene glycol (Figures S1, 2, 3). The changes in the chemical and physical properties of the polymer when incubated with BP8, and the generation of diverse degradation products, some of them potential metabolic intermediates in the degradation process, are evidences of the BP8’s degradative capability, which is sustained by the diverse xenobiotic degrading enzymes encoded in its metagenome (Table 1). Some of the biodegradation products (Figure 2) seem to be the result of oxidative reactions on C-C bonds flanking TDI, MDI or the acrylates’ styrene ring (Figure 3, S1), generating aromatic compounds containing hydroxyl, aldehydes or organic acids. Additionally, the copolymer aromatic compounds could be destabilized by monooxygenases, which introduces hydroxyl groups to the aromatic rings, and by dioxygenases that catalyzes reductive dihydroxylation, generating central intermediates that can be cleaved by dearomatizing dioxygenases producing carboxylic acids [73, 74]. The enzymes for the complete benzoate metabolism are encoded in the BP8 metagenome and could account for PE-PU-A aromatic rings catabolism (Table 1). Aliphatic chains from acrylates and polypropylene glycols can be metabolized by alkane 1-monooxygenases, that activate aliphatic chains by terminal or subterminal oxidations and by the activities of ADH and ALDH, generating compounds that can be channeled by beta-oxidation into the fatty acids metabolism (Table 1). If terminal oxidations are introduced, primary alcohols are generated and transformed into aldehydes, carboxylic acids and acyl-CoA. If subterminal oxidations of aliphatic chains occur, secondary alcohols are formed, which upon breakdown, will produce ketones and thereafter esters, which are hydrolyzed to alcohol and organic acids [75, 76]. Many different esters compounds were identified in the BP8’s degradation products, suggesting that subterminal oxidation of alkanes could be an important route in PU metabolism (Figures 2, 3, S1). The cleavage of ester bonds by PU-esterases would produce alcohols and organic acids, and the cleavage of urethane groups by carbamate-hydrolases would produce nitrogen-containing compounds and aromatic isocyanate derivatives. As we detected these degradation products by GC-MS analysis (Table 1, 2, Figure 2), hydrolysis of ester and urethane bonds are accomplished during PE-PU-A degradation by BP8. The identification of several PU-esterases and carbamate hydrolases encoded in most of the BP8 genomes support this conclusion (Table 2).

The metabolic reactions proposed for the degradation of the additives and the PE-PU-A copolymer present in PolyLack^®^ by the BP8 community are based on the phenotypic potential encoded in its metagenome. The use of proximity ligation Hi-C technology allowed to define, with high confidence, what genes belong to each of the different species of BP8 (Table 1). In this community, xenobiotic degradation is a niche dominated by *Paracoccus* sp. BP8 and *Ochrobactrum intermedium* BP8.5, in whose genomes, key enzymes for different steps of biodegradation are widely represented (Table 1), which must be the reason for their preponderance in the BP8 community. In addition, Microbacteriaceae bacterium BP8.4 genome encodes enzymes for the metabolism of aromatic compounds suggesting that metacleavage ring excision and muconate lactone formation might be functional. On the other hand, *Chryseobacterium* sp. BP8.2 and *Parapedobacter* sp. BP8.3 genomes, harbor genes encoding complementary metabolic activities for alkanes oxidation, such as hydrolysis and oxidation of linear intermediates. The finding of such a diverse genetic repertoire in the BP8 metagenome suggests a remarkable metabolic versatility, with strong hydrolytic and oxidative capabilities that can play significant roles in the degradation of diverse environmental contaminants. The abundance and distribution of these catabolic enzymes among the different members of the BP8 community, suggest syntrophic mechanisms driving community behavior. However, incomplete genome reconstruction in the deconvolution analysis, resulting in potential pathway gaps in certain genomes, cannot be ruled out, nor can the collapsing of multiple strains into a single cluster. On the other hand, although *Paracoccus* and *Ochrobactrum* are predominant in the BP8 community by far, we cannot discard that specific enzymatic activities encoded in genomes of little abundant species can be crucial for the successful performance of BP8.

The present work provides deep understanding of the microbial ecology of a selected landfill microbial community capable of PU and xenobiotics degradation by revealing its composition and its outstanding phenotypic potential observed in the catalytic capabilities that its members could display to cleave different recalcitrant functional groups. Altogether, these features place BP8 community as a quite promising source for developing environmental biotechnology strategies contributing to mitigate anthropogenic plastics and xenobiotics pollution for achieving better environmental quality. Moreover, further exploration of individual species of the community will allow the manipulation of novel catabolic capabilities in order to improve biodegradative technological processes.

## ACKNOWLEDGEMENTS

IG and MB acknowledge Consejo Nacional de Ciencia y Tecnología for their Ph.D. scholarships. ASR acknowledges Dirección General de Asuntos del Personal Académico, UNAM, for his scholarship for a posdoctoral position at Facultad de Química, UNAM. We thank USAII-FQ analytical support provided by Ch. Marisela Gutiérrez Franco, Ch. Rafael Iván Puente Lee, Ch. Víctor Hugo Lemus and Ch. Elvira del Socorro Reynosa in the FTIR, SEM, carbon quantification, and thermogravimetry analyses, respectively. The technical support of MSc. Fernando de Jesús Rosas Ramírez in the use of the GC-MS equipment is appreciated. Also, we are grateful to Gerardo Cedillo, Salvador López and Karla E. Reyes from Institute of Materials Research (IIM-UNAM) for their assistance in NMR, GPC and DSC analyses, respectively, and to Taylor Reiter and C. Titus Brown for help with metagenomic analysis.

## FUNDING INFORMATION

This work was funded by Programa de Apoyo a Proyectos de Investigación e Innovación Tecnológica, Dirección General de Asuntos del Personal Académico, Universidad Nacional Autónoma de México grants IN217114 and IN223317, and Programa de Apoyo a la Investigación y el Posgrado, Facultad de Química, Universidad Nacional Autónoma de México, grant 5000-9117. IL, MP, and SS were supported in part by NIAID grant 2R44AI122654-02A1.

## CONFLICT OF INTEREST STATEMENT

IL, MP, and SS are employees and shareholders of Phase Genomics, a company commercializing proximity ligation technology.

---

**Table S1.** Effects of BP8 biodegradative activity on the PE-PU-A copolymer analyzed by Differential Scanning Calorimetry.

| **Culture time (days)** | **Tg (°C)** | **Tm-I (°C)** | **Tm-II (°C)** | **Tm-III (°C)** | **Tc (°C)** |
| --- | --- | --- | --- | --- | --- |
| Non-inoculated | 50.2 | 70.0 | 210.6 | 398.1 | 459.6 |
| 5 | 39.5 | 68.8 | 211.0 | 408.7 | 478.2 |
| 10 | 46.0 | 68.0 | 210.1 | 407.9 | 480.2 |
| 15 | 38.1 | 70.5 | 211.1 | 393.2 | 479.9 |
| 20 | 46.2 | 78.8 | 213.9 | 392.5 | 476.2 |

Tg = glass transition temperature; Tm = melting temperature; Tc = crystallization temperature.

**Table S2.** Molecular weight and polydispersity index of the PE-PU-A copolymer during cultivation with the BP8 community.

| **Culture time (days)** | ***Mn***<br>**(g/mol)** | **PDI** |
| --- | --- | --- |
| Non-inoculated | 101 896 ± 8098 | 2.0 ± 0.3 |
| 5 | 96 798 ± 712 | 2.2 ± 0.1 |
| 10 | 89 258 ± 5825 | 2.5 ± 0.3 |
| 15 | 82 577 ± 1168 | 3.1 ± 0.6 |
| 20 | 79 859 ± 9225 | 2.4 ± 0.3 |
| 25 | 65 609 ± 990 | 2.6 ± 0.2 |

Number-average molecular weight (*Mn*) and polydispersity index (PDI) were calculated by Gel Permeation Chromatography with monodisperse polystyrene calibration standards and using THF as eluent. n=3

**Table S3.** General features of the deconvoluted genomes from the BP8 metagenome.

| Cluster ID | Genome Size (bp) | Num Contigs | Contig N50 | <sup>a</sup>Completeness (%) | <sup>b</sup>Relative Abundance (%) | Novelty Score (%) | GC (%) | Identification | Genes assigned | Proteins assigned |
| --- | --- | --- | --- | --- | --- | --- | --- | --- | --- | --- |
| 1 | 4 275 656 | 282 | 51 004 | 89.4 | 57.7 | 1.6 | 67.8 | *Paracoccus* sp. BP8 | 4 225 | 4 073 |
| 2 | 2 157 639 | 388 | 7 081 | 95.6 | 3.7 | 98.7 | 47.3 | sp. BP8.2 | 2 253 | 185 |
| 3 | 5 478 545 | 1098 | 6 493 | 95.5 | 12.5 | 99.2 | 48.1 | *Parapedobacter* sp. BP8.3 | 5 310 | 5 173 |
| 4 | 2 790 120 | 158 | 39 967 | 97.7 | 3.6 | 94.0 | 71.3 | bacterium BP8.4 | 2 850 | 705 |
| 5 | 2 916 513 | 1146 | 2 823 | 71.0 | 22.5 | 2.5 | 58.4 | *Ochrobactrum intermedium* BP8.5 | 3 472 | 3 162 |

<sup>a</sup>Completeness was calculated based on 40 single copy gene markers [36]. All the genomes' drafts have at least 18 tRNAs and, except for cluster 5, at least 1 rDNA gene copy.
<sup>b</sup>Relative abundance was normalized according to the reads distribution along the deconvoluted taxon.
<sup>c</sup>For Microbacteriaceae bacterium BP8.4 no further classification was possible even that nine single-copy markers were analyzed.

**Table S4.** ^a^Phylogenetic relatedness of the bacterial species from the BP8 community identified by Hi-C metagenome deconvolution.

| Clusters/<br>Classification/<br>GeneBank Acc.<br>Num. | Organism | Assembly | Genome<br>size<br>(Mb) | GC<br>content<br>(%) | <sup>a</sup>ANI<br>value<br>(%) | Proteins<br>encoded<br>in the<br>genomes |
| --- | --- | --- | --- | --- | --- | --- |
| Cluster 1/<br>*Paracoccus* sp.<br>BP8<br>GCA_003852815.1 | *Paracoccus* sp. J39 | GCA_000518925.1 | 4.42837 | 68.08 | 98.73 | 4131-4993 |
|  | *Paracoccus denitrificans* | GCA_000203895.1 | 5.23619 | 66.78 | 90.24 |  |
|  | *Paracoccus aminovorans* | GCA_001546115.1 | 4.58940 | 67.50 | 85.59 |  |
|  | *Paracoccus halophilus* | GCA_900111785.1 | 4.00871 | 65.20 | 82.17 |  |
|  | *Paracoccus yeei* | GCA_002073635.2 | 4.82967 | 67.08 | 81.95 |  |
| Cluster 2/<br>*Chryseobacterium*<br>sp. BP8.2<br>GCA_003852805.1 | *Chryseobacterium koreense* | GCA_001045435.1 | 3.15420 | 40.10 | 69.38 | 3000-4810 |
|  | *Chryseobacterium camelliae* | GCA_002770595.1 | 4.37635 | 41.80 | 68.86 |  |
|  | *Chryseobacterium luteum* | GCA_000737785.1 | 4.71855 | 37.30 | 68.70 |  |
|  | *Chryseobacterium* sp. 52 | GCA_002754245.1 | 5.29882 | 37.00 | 68.49 |  |
|  | *Chryseobacterium antarcticum* | GCA_000729985.1 | 3.12366 | 36.10 | 68.10 |  |
| Cluster 3/<br>*Parapedobacter*<br>sp. BP8.3<br>GCA_003852785.1 | *Parapedobacter indicus* | GCA_900113765.1 | 6.15523 | 48.00 | 80.23 | 3796-4949 |
|  | *Parapedobacter koreensis* | GCA_900109365.1 | 5.54776 | 48.20 | 74.24 |  |
|  | *Parapedobacter luteus* | GCA_900168055.1 | 4.82992 | 49.30 | 73.66 |  |
|  | *Parapedobacter composti* | GCA_900112315.1 | 4.62202 | 50.00 | 73.32 |  |
| Cluster 4/<br>Microbacteriaceae<br>bacterium BP8.4<br>GCA_003852775.1 | *Micrococcales bacterium* 72-143 | GCA_001898835.1 | 3.33048 | 71.60 | 85.07 | 2535-3047 |
|  | <sup>b</sup>*Leifsonia* sp. Leaf336 | GCA_001423695.1 | 4.15779 | 69.60 | 74.40 |  |
|  | <sup>b</sup>*Microcella alkaliphila* | GCA_002355395.1 | 2.70284 | 68.40 | 74.10 |  |
|  | <sup>b</sup>*Plantibacter* sp. H53 | GCA_001650455.1 | 4.01278 | 69.40 | 73.98 |  |
|  | <sup>b</sup>*Herbiconiux* sp. YR403 | GCA_000799285.1 | 3.60404 | 62.00 | 71.66 |  |
|  | <sup>c</sup>*Arthrobacter cupressi* | GCA_900099975.1 | 4.0588 | 67.00 | 70.82 |  |
| Cluster 5/<br>*Ochrobactrum*<br>*intermedium* BP8.5<br>GCA_003852825.1 | *Ochrobactrum intermedium*<br>LMG 3301 | GCA_000182645.1 | 4.72539 | 57.70 | 98.48 | 3645-4502 |
|  | *Ochrobactrum anthropi* | GCA_000017405.1 | 5.20578 | 56.15 | 87.48 |  |
|  | *Ochrobactrum oryzae* OA447 | GCA_002943495.1 | 4.46700 | 56.20 | 87.19 |  |
|  | *Brucella ceti* | GCA_000590795.1 | 3.27803 | 57.27 | 82.03 |  |
|  | *Ochrobactrum* sp. P6BS-III | GCA_002016635.1 | 5.25313 | 56.00 | 81.19 |  |

<sup>a</sup> Phylogenetic relatedness was calculated based on ANI value with OrthoANLu algorithm. ANI values higher than 95% indicate that the compared genomes are the same species [39].
<sup>b</sup> Members of Microbacteriaceae family, order Micrococcales.
<sup>c</sup> Member of Micrococcaceae family, order Micrococcales.

**Figure S1.**
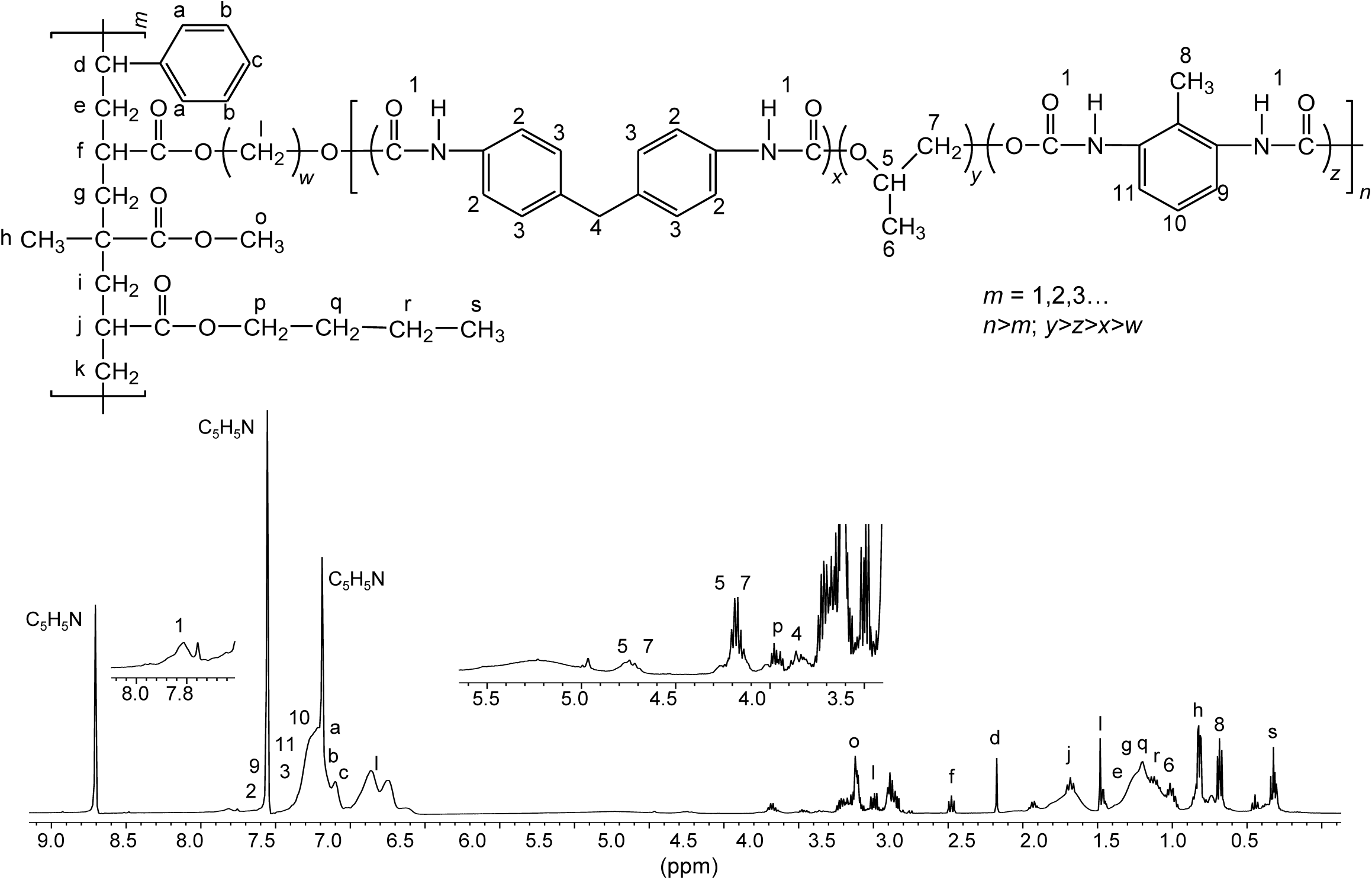
Proposed chemical structure for the PE-PU-A copolymer present in PolyLack^®^. This structure was proposed based on the ^1^H-NMR analysis of dried PolyLack^®^, the information included in the manufacturer technical manual [SayerLack. Poly Lack Aqua Mate UB-0810. Manual Técnico. 2013. http://www.gruposayer.com//web/uploads/file/TDS_UB-0810-file160515563.pdf], the GC-MS analysis (Figure 2), and the most frequent acrylates used in the synthesis of these types of copolymers [Maurya SD, Kurmvanshi SK, Mohanty S, Nayak SK. A review on acrylate-terminated urethane oligomers and polymers: synthesis and applications. Polym Plast Technol Eng. 2018; 57:625-56; Pardini OR, Amalvy JI. FTIR, 1H-NMR spectra, and thermal characterization of water-based polyurethane/acrylic hybrids. J Appl Polym Sci. 2008; 107:1207-14]. Synthesis of PE-PU-A copolymers starts by the polycondensation of polyols (polypropylene glycol) (y moiety) and diisocyanates (TDI and MDI) (x and z moieties) followed by end capping with acrylates’ mixture (m moiety). From the most frequently used acrylates we selected methyl methacrylate, butyl acrylate, hydroxy acrylate and styrene as representatives in this structure. In the ^1^H-NMR spectrum, chemical shifts are provided in parts per million from SiMe_4_ as internal reference. Signal 1 is assigned to carbamate groups (NH-COO); signals a, b, c, 2, 3, 9-11 are assigned to the aromatic protons; signals 4 and 8 correspond to the protons of methylene (CH_2_) and methyl (CH_3_) groups in MDI and TDI, respectively; signals 5-7 correspond to PPG; signals l correspond to the hydroxyl proton (CH_2_-O) and methylene groups (CH_2_) in the chain of hydroxy-acrylate; signals f, j, o and p correspond to the acrylic groups (CH-COO, CH_2_-COO or CH_3_-COO), signals d (CH), e, g, i, k, q, r (CH_2_), h and s (CH_3_) are assigned to methylene and methyl groups in the acrylate mixtures.

**Figure S2.**
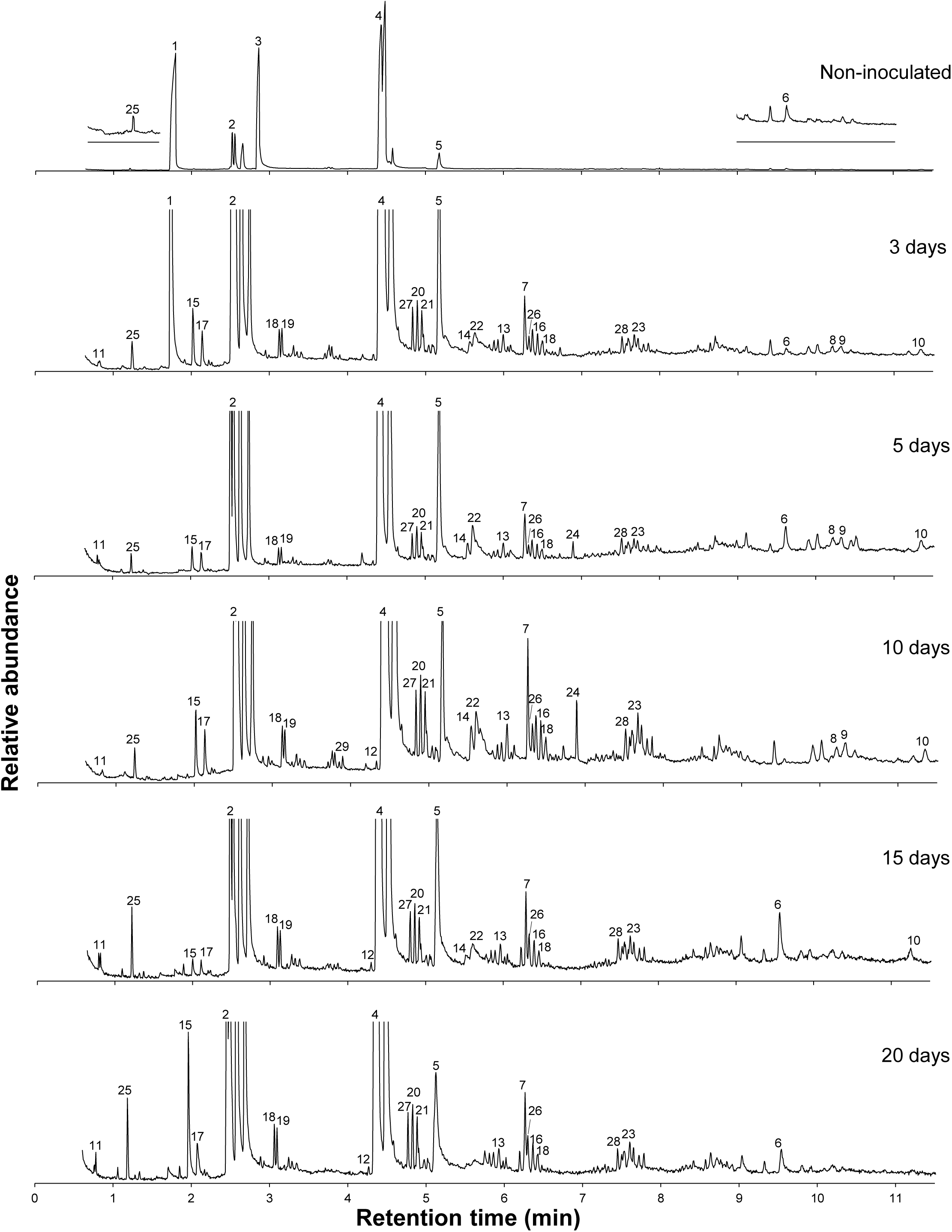
Chromatograms of cell-free supernatants from BP8 cultures grown in MM-PolyLack. Numbered signals designate compounds identified by mass spectrometry listed in Figure 2. Chromatogram of non-inoculated sample is at original scale, whereas the other chromatograms are at larger scale.

**Figure S3.**
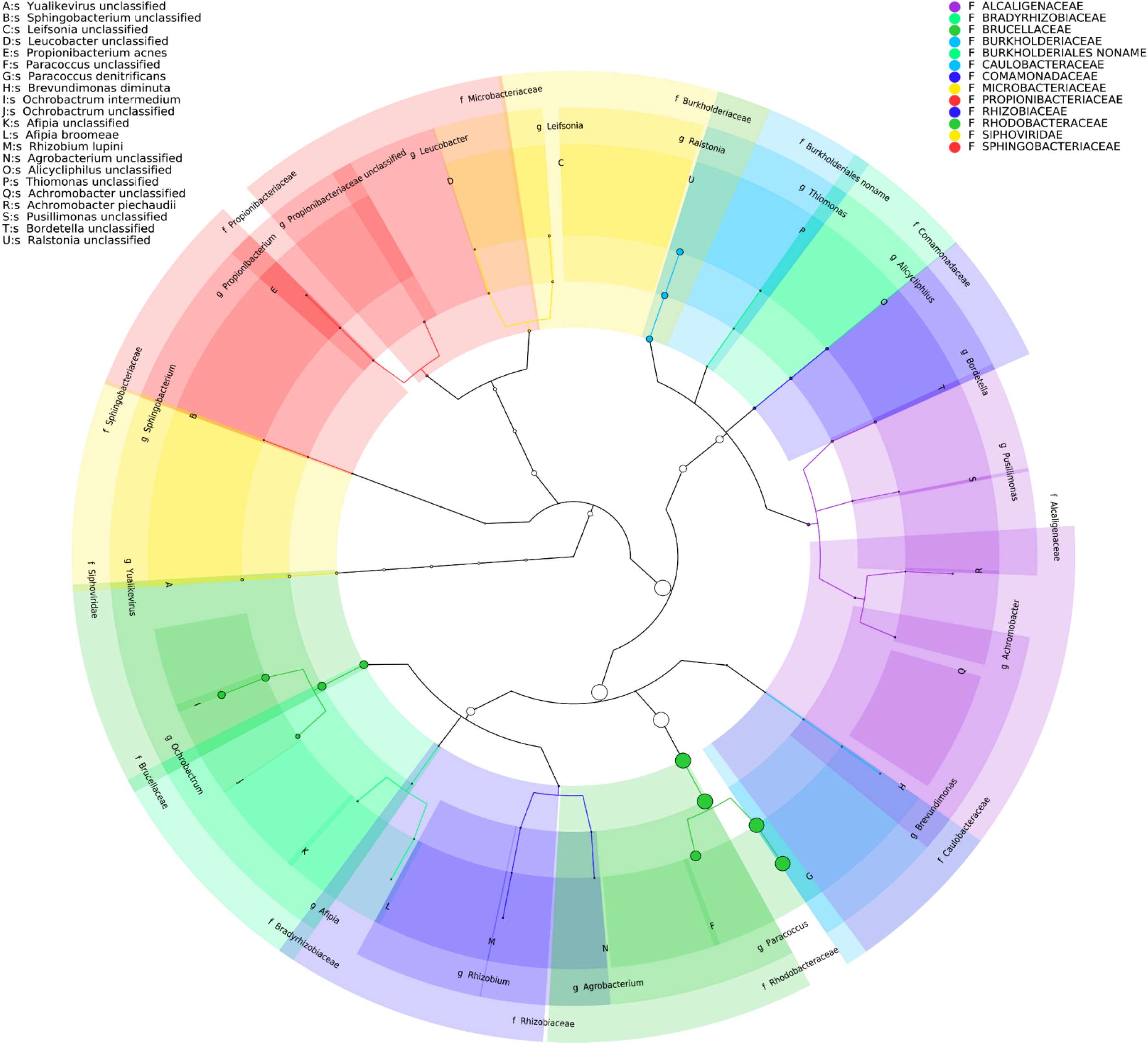
Taxonomic cladogram of BP8 community microbial diversity profiled with MetaPhlAn. Circles size is proportional to the taxon relative abundance. The most abundant taxa were *Paracoccus* genus (83.3%) and *Ochrobactrum* genus (8.7%). Families are color-labeled and predicted species diversity is indicated by capital letters [Asnicar F, Weingart G, Tickle TL, Huttenhower C, Segata N. Compact graphical representation of phylogenetic data and metadata with GraPhlAn. PeerJ. 2015;3:e1029].

**Figure S4.**
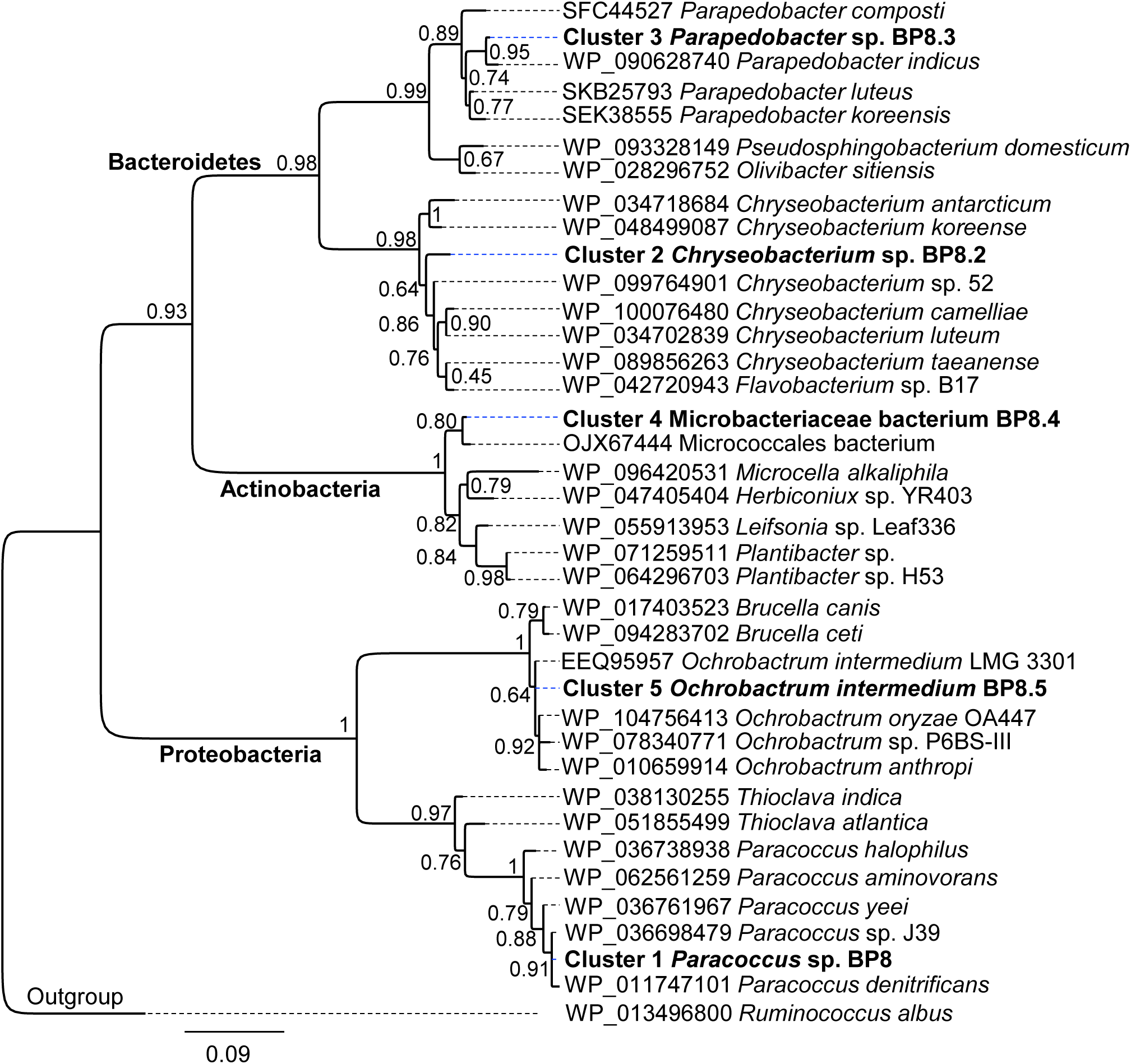
Maximum likelihood phylogeny for taxonomic delimitation of the deconvoluted genomes from the BP8 metagenome. This analysis was performed with three phylogenetic markers, ribosomal protein L3, ribosomal protein L5 and DNA gyrase A subunit, which generated similar results. Here we present the analysis for ribosomal protein L3. Branch support values are indicated in the corresponding nodes. Bar indicates the number of expected substitutions per site under the WAG + G model. A sequence of *Ruminococcus albus* (Firmicutes) was used as outgroup. Key genome clusters are highlighted in bold and different Phyla are indicated at the left. Sequences for L3 ribosomal proteins of the deconvoluted genomes are accessible in the NCBI GenBank under accession numbers RQP07704.1, RQP15098.1, RQP16503.1, RQP08603.1 and RQP16393.1 for clusters 1 to 5, respectively.

## Notes

#### Summary of Updates

This version of the manuscript has been revised to update the Supplementary material

